# Inferring Migration Networks with Time-Lagged *F*_2_ Statistics

**DOI:** 10.64898/2026.03.12.710875

**Authors:** Giulio Isacchini, Takashi Okada, Clemens Schmid, Divyaratan Popli, Benjamin M Peter, Stephan Schiffels, Oskar Hallatschek

## Abstract

Major demographic events, such as population bottlenecks, founder effects, range expansions, and admixture events, have left lasting imprints on human genetic diversity. Ancient DNA (aDNA) sequencing now makes it increasingly possible to observe these signals across time, opening new avenues to address long-standing questions in human demographic history. Yet, deciphering this genetic archive of demographic history is challenging due to high levels of noise, and the complex ways in which demographic processes shape genetic variation. Here, leveraging the linear time evolution of the expectation of neutral allele frequencies, we develop a method to uncover systematic patterns of gene flow from metapopulation time-series data. We show that directional migration rates can be inferred via linear regression on time-dependent genetic dissimilarity between populations, quantified by an extended *F*_2_ statistic evaluated between successive time points. Despite small sample sizes, the method reliably infers migration rates from simulated data by integrating information across multiple time slices. Applied to aDNA sampled from the last 6000 years, we recover signals of well-documented migrations and infer an ancient pan-European migration network. While complementing existing tools that estimate static ancestry proportions, our framework tracks how ancestry is dynamically redistributed through time.

## Introduction

Ancient genomes preserve a noisy record of past migrations, expansions, replacements, and extinctions—events once thought irretrievable but now increasingly accessible through ancient DNA (aDNA) analysis. Over the past decade, extensive work has reconstructed aspects of human demographic history from such data (1–7). Yet most studies have focused on specific spatial or temporal contexts or on time-aggregated signals, such as ancestry proportions inferred from present-day or temporally averaged populations (8–12). How mobility patterns evolve continuously through time remains poorly understood, in part because sparse and uneven genome sampling limits statistical power, and the high levels of missing data typical in aDNA complicate cross-sample comparisons.

A recent study (13) showed that allele frequency changes over the past 5,000 years can be explained by gene flow alone, without invoking detectable natural selection. Motivated by this result, we develop a method to infer temporal patterns of gene flow under a neutral evolutionary model, using allele frequency time series. Our framework operates directly on genomic data and avoids assumptions about the existence or composition of ancestral source populations that underlie admixture proportion estimates (14).

Previous efforts have mapped effective migration surfaces from genetic data (2, 4, 15), including large-scale applications to human diversity (5), but these methods omit temporal information and cannot resolve changes in migration over time. Spatio-temporal interpolation of genetic profiles helps resolve individual-level migration events (16), with limited quantitative interpretation at the population-level. Alternatively, continuous diffusive spread has been modeled via partial differential equations (17), a formulation that struggles to capture discrete, long-distance migration events.

We exploit the fact that gene-flow impacts the covariance of neutral allele-frequency changes between populations over time. Without interaction, allele frequency changes should be uncorrelated, resulting in zero covariance. At the other extreme, if two populations are highly connected - effectively behaving as a single mixed population - their allele frequencies will be strongly correlated (Fig. 1). In between these two extremes, our method seeks to infer a gene flow matrix **A**_*ij*_ by measuring the covariance of allele frequencies across different regions.

**Fig. 1.**
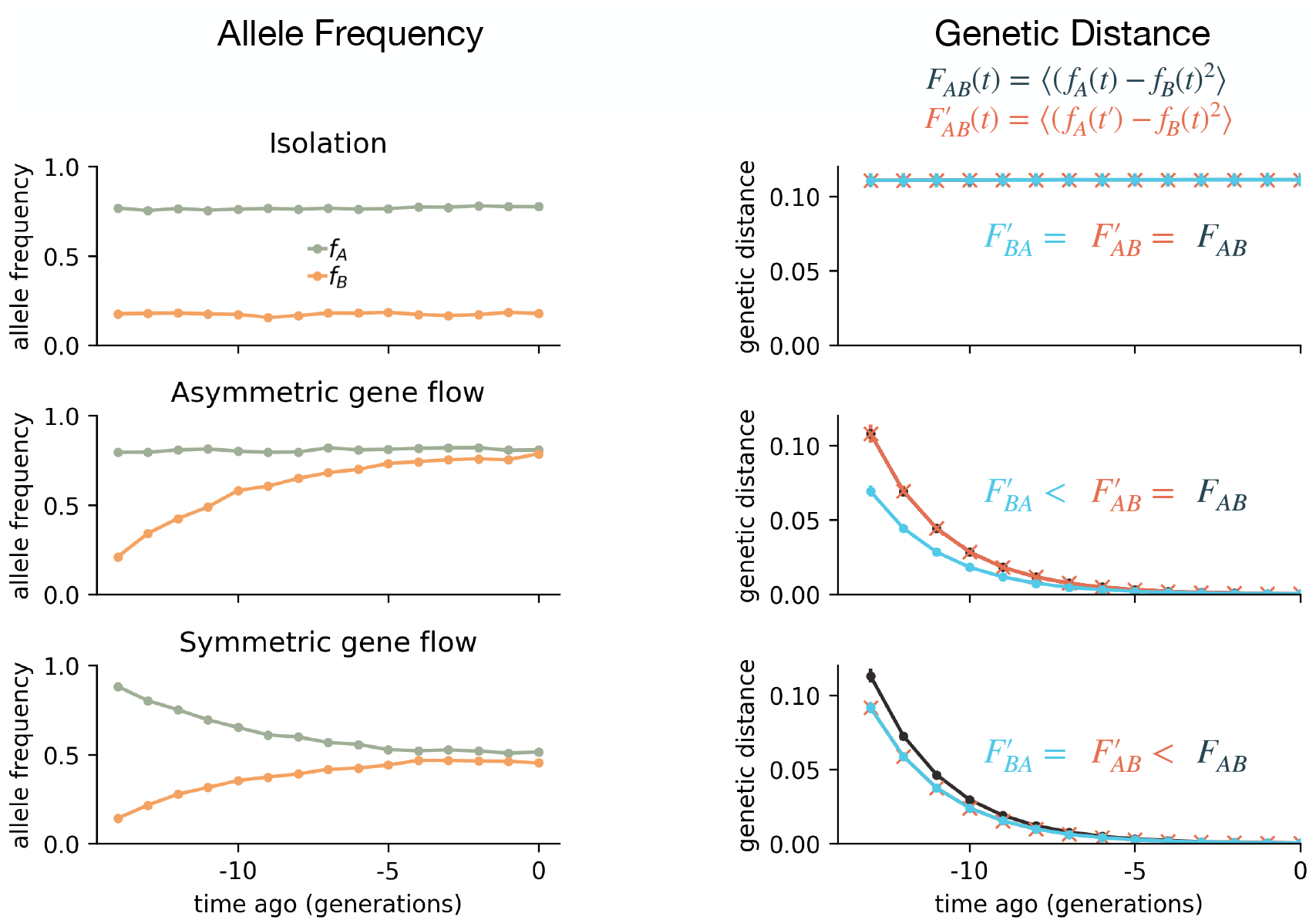
A decline in genetic distance between populations signals gene flow. In ancient DNA research, genetic distance is often measured as the mean squared allele frequency difference between populations, known as the *F* -statistic. Here, we introduce a time-lagged extension, *F* ^*′*^, which quantifies the genetic distance between populations sampled at different time points (see main text for definitions). We illustrate how directional gene flow is reflected in the temporal decay of *F* and *F* ^*′*^ by simulating allele frequency trajectories for two populations, A and B, under three scenarios: no migration (top), strong unidirectional migration (middle), and symmetric migration (bottom). All alleles were initialized with *f*_*A*_ = 0.8 and *f*_*B*_ = 0.2. Migration reduces pre-existing allele frequency differences (left panels), leading to a decay in all genetic distance measures (right panels). Comparing *F* and *F* ^*′*^ reveals directionality: with unidirectional flow from A to B (middle), 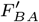 declines faster than *F*_*AB*_ and 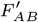 (right middle panel), whereas symmetric migration produces no such asymmetry (bottom panels). Simulation details are provided in the SI. Default parameters: *σ* = 1, *T* = 15, *l* = −1, *N*_*D*_ = 2, *N*_traj_ = 5000, *N*_pop_ = 1000, *N*_*S*_ = 5000. Migration matrices: **A** = *I* for isolation, **A** = [(1, 0), (0.2, 0.8)] for asymmetric flow, and **A** = [(0.9, 0.1), (0.1, 0.9)] for symmetric flow. Only one representative lineage pair is shown.

This general approach has already proven effective in reconstructing transmission networks during the Covid-19 pandemic (18, 19), where SARS-CoV-2 lineage frequency data was used to estimate inter-regional importation rates using a Kalman Filter that accounted for sampling noise and genetic drift under Gaussian assumptions (20). While such assumptions were appropriate given the dense sequencing data, they are less applicable to ancient DNA (aDNA), where sparse sampling results in many regions (currently) lacking data for extended periods. Nonetheless, the larger human genome and the presence of recombination yield a far greater number of approximately independent lineages compared to the SARS-CoV-2 dataset, potentially enabling inference even with far fewer sampled genomes.

To harness this abundance of lineages, we aggregate information across genomes to quantify allele frequency covariance between populations. These covariances are expressed through the *F*_2_ statistics, which measure the genetic distance between populations (21, 22). Widely used in aDNA studies (23, 24), *F*_2_ statistics are typically computed pairwise across populations as a preprocessing step for dimensionality reduction via multidimensional scaling (25). Their empirical estimators naturally incorporate binomial sampling noise, and their calculation using block-bootstrap methods is robust to linkage disequilibrium (26).

We show that the *F*_2_-distance between populations sampled at different times, which we call time-lagged 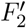 statistics, is linearly related to the inter-population genetic distances between populations sampled at the same time. The linear coefficients characterize gene flow and can be efficiently inferred via a maximum likelihood approach. We estimate confidence intervals by taking into account uncertainty and potential biases due to sparse sampling, temporal dating and genetic linkage. The strength of this method lies in its efficiency and scalability. As the efforts in sequencing will continue to expand the pool of aDNA samples available, our method will be able to extend to longer time scales than those studied in the present work.

## Results

### Model and Inference

Genetic dissimilarity between populations can be robustly quantified via *F*_2_-statistics (21, 22). Given two populations *A* and *B* sampled at time *t*, the *F*_2_ statistic is defined as *F*_*AB*_(*t*) = 𝔼 [(*f*_*A*_(*t*) − *f*_*B*_(*t*))^2^] where the expectation is taken over all sites in the genome, and *f*_*K*_ indicates the frequency of the allele at the site in population *K*. Typically, analyses involve comparing predetermined populations in an all-to-all fashion, without considering the dating of the samples.

Here, we study how *F*_2_-statistics change over time and demonstrate that it is possible to deduce directional gene flow from these temporal changes. We introduce an additional quantity, that we refer to as time-delayed F-statistics and it is defined as 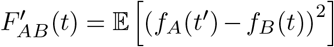 where *t*^*′*^ = *t* + Δ*t* is delayed by a factor Δ*t*. Fig. 1 illustrates how the strength, direction, and asymmetry of gene flow each imprint characteristic signatures on the statistics *F*_*AB*_, 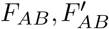, and *F*_*BA*_ - the key signal leveraged by our inference framework.

We model the dynamics of neutral alleles in a population composed of *n* sub-populations as a function of genetic drift and migration, similarly to (18). Due to the sample sparsity of aDNA, accurate estimation of single allele frequencies is challenging. For this reason, we rewrite the dynamics in terms of *F*_2_ summary statistics, which can be robustly estimated (22). In Methods, we show that the temporal change in *F*_2_ statistics is given by

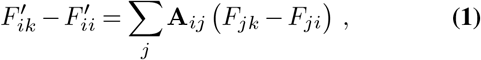

where **A**_*ij*_ is an *n* × *n* migration matrix: its ele nts are non-negative and, within each row, sum up to one, ∑_*j*_ **A**_*ij*_ = 1. The coefficient **A**_*ij*_ represents the proportion of individuals in population *i* that are replaced by migrants from population *j* within the considered time period, thus capturing migration rates between regions. **A**_*ij*_ can also be interpreted as a matrix of backward-in-time probabilities that a lineage moves from population *i* to population *j* in the time period Δ*t*.

The linearity of Eq. 1 implies that the migration matrix **A**_*ij*_ can be inferred via linear regression. In constructing confidence intervals, we take into account uncertainty and potential biases due to random sampling of individuals, uncertainty in the temporal assignment, and genetic linkage across loci, see Fig. S1 and Methods for additional details.

### Simulation-based validation of inference method

We study the applicability of our method to sparse, incomplete aDNA data in a simulated scenario. We perform simulations using msprime (27, 28) and population parameter specifications from the stdpopsim project (29, 30) for a diploid human population with a single chromosome with uniform recombination rate. As shown in Fig. 2A, we simulate a scenario where an ancestral human population of size *N*_*e*_ = 104 splits into 4 deeply diverged populations, which begin to expand exponentially for 5000 generations. Migration is defined via a random migration matrix **A**_true_ whose rows sum up to 1 and begin 50 generations before the end of the simulations. At the same time, we begin sampling 20 individuals from each population at intervals of 10 generations apart. Given the approximate generation time of humans is ∼30*y*, this reproduces sampling times of Δ*t* = 300*y*, similar to those in the aDNA data sets analyzed below. We sample one chromosome per individual and introduce missing data at 20% probability, to reproduce the sampling noise of typical pseudohaploid aDNA data. Fig. 2B presents an example of the neutral allele frequency trajectories, which serve as the input for our approach. Fig. 2C illustrates the temporal variation of the *F* statistics during the sampling period. We refer to Fig. S2, for an overview of how genetic divergence gets established in the time period when the populations are isolated.

**Fig. 2.**
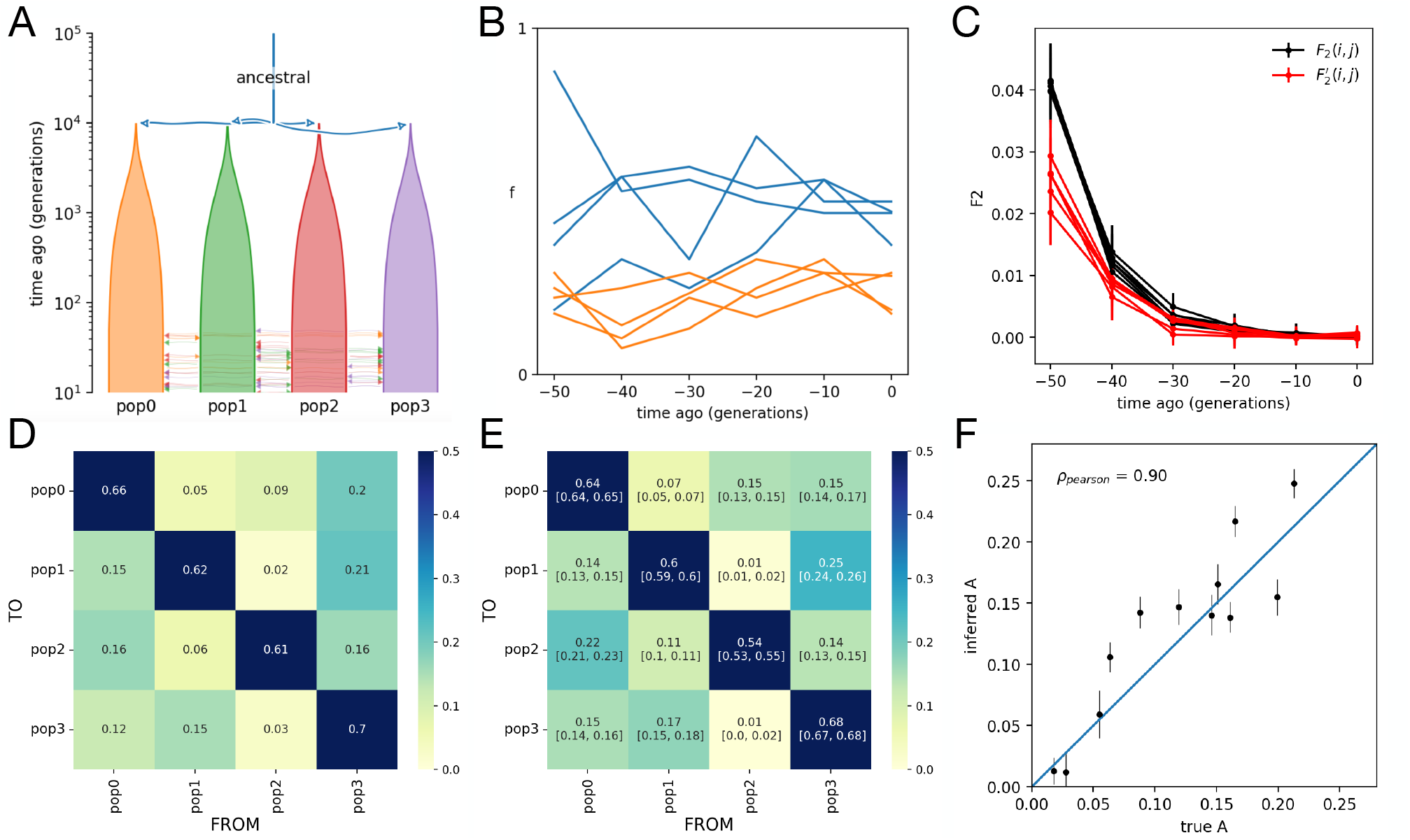
Simulations to assess inference on sparse, incomplete data. **(A)** Four diploid human populations are simulated as descendants of a single ancestral population of size *N* = 10^4^. A single chromosome with a uniform recombination rate is simulated using standard `stdpopsim` parameters, with 20% of data randomly removed. Initial isolation and exponential expansion increase divergence between populations. Migration, defined by a random matrix **A**, begins 100 generations before the present. Sampling starts at this time and is repeated every 10 generations, matching the time discretization of the empirical analysis. **(B)** Example allele frequency trajectories for two loci (blue and orange); each line of the same colour corresponds to a different population. **(C)** Genetic divergence (*F*_2_ and 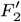 statistics) decreases during the migration period. **(D)** Reference migration matrix used in the simulation. **(E)** Inferred migration matrix with confidence intervals given by lower and upper quartiles. **(F)** The inferred matrix closely matches the reference; the Pearson correlation between off-diagonal true and average inferred coefficients is *ρ*_Pearson_ = 0.9.

In our simulation scenario, migration significantly reduces the genetic divergence between populations — divergence that initially emerged through genetic drift and population expansion. The correlation plot in Fig. 2F demonstrates strong agreement between inferred and true migration coefficients. An extended benchmark across a broad range of parameters using Wright–Fisher simulations is presented in Fig. S3.

### Early Neolithic expansion

As an initial application to real data, we examine the early Neolithic expansion—a well-documented migration from the Levant into Europe supported by both archaeological (31–33) and archaeogenetic evidence (34–37). For this analysis, we selected ancient genomic samples dated between 6800 and 5600 BCE from Southeastern Europe (SE Europe) and the Levant/Anatolia region (LevA) (see Methods for details on the data preparation (16, 38, 39)). We organized these samples into two distinct populations and established time intervals of 200 years each, as shown in Fig. 3A. We calculated *F*_2_-statistics and time-delayed 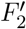-statistics for all possible combinations of these populations.

**Fig. 3.**
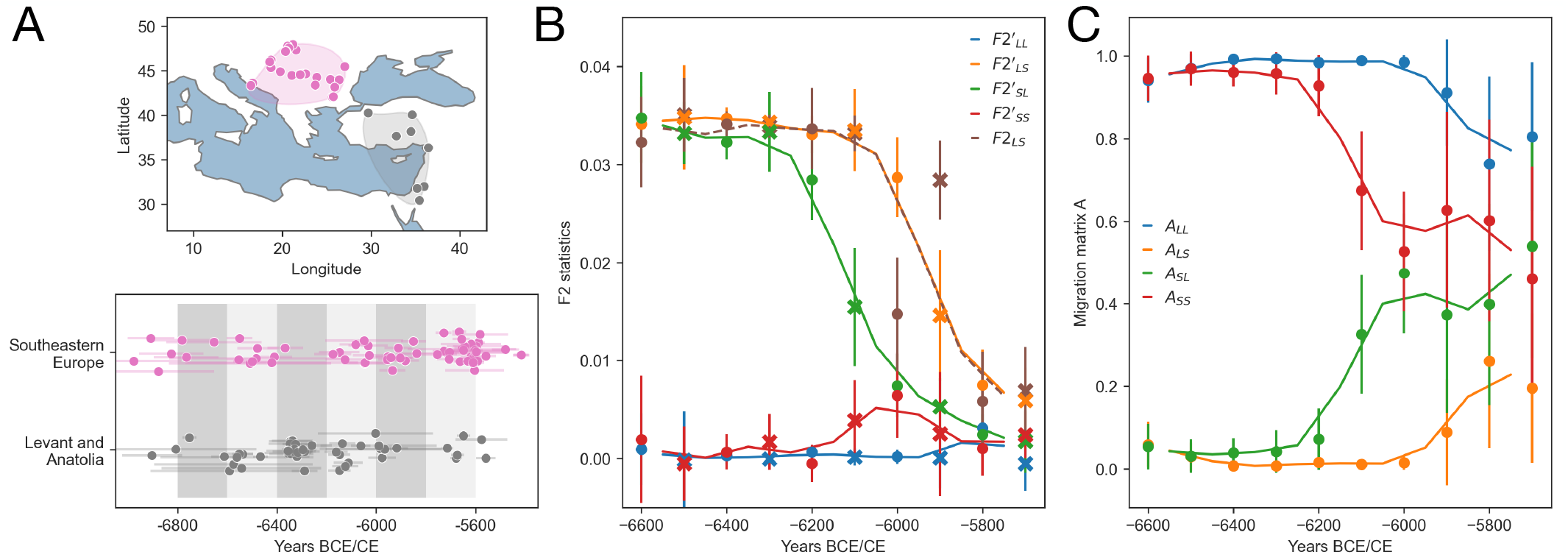
Early Neolithic expansion through time. **(A)** Spatio-temporal distribution of the samples analyzed, focusing on the Early Neolithic expansion from the Levant and Anatolia into Southeastern Europe. Time windows of Δ*t* = 200 y from 6800 to 5600 BCE yield six time steps, shown in shades of gray. **(B)** *F*_2_ distances across time windows. The solid line identifies the average between the values computed between the six time steps and the corresponding values with respect to the overlapping delayed time windows computed at *dt* = 100*y*. **(C)** Inferred migration across time windows. As in the previous figure, the solid line represents the average across overlapping delayed time windows.

In this simple scenario with just two populations, the patterns present in the *F* -statistics are directly represented in the inferred migration rates (Fig. 3C): The inferred migration rate from LevA to SE Europe (indicated by the green line) rises significantly above zero, while the coefficient from SE Europe to LevA (represented by the orange line) remains close to zero throughout the entire period. This observation aligns with the presence of substantial gene flow from the Levant to Southeastern Europe, consistent with the archaeological record as discussed in (40–42). For an estimation of the relaxation times, see Fig. S5.

The analysis indicates that observing the temporal dynamics of *F*_2_-statistics can be helpful in characterizing the source of change in the genetic composition of a population and that our method can effectively identify known migration patterns.

### Reconstructing Gene Flow Dynamics Across Nine Regions

In the previous section, we tested the validity of our method in a limited and well-understood spatio-temporal context. We now seek to characterize the change in the genetic composition of nine populations spanning a duration of 3000 years, ranging from 4000 to 1000 BCE, see Fig. 4A-B for the spatiotemporal distribution of the samples. In this time period, linguistic, archaeological and – most recently – genetic evidence hints towards a major migration from the Pontic steppes into Central Europe, potentially corresponding to the spread of the Corded Ware culture and Indo-European languages into Central Europe (8, 9, 12). Incidentally, the significance and magnitude of this event had not been appreciated by archaeologists prior to the insights gained through ancient DNA (43).

**Fig. 4.**
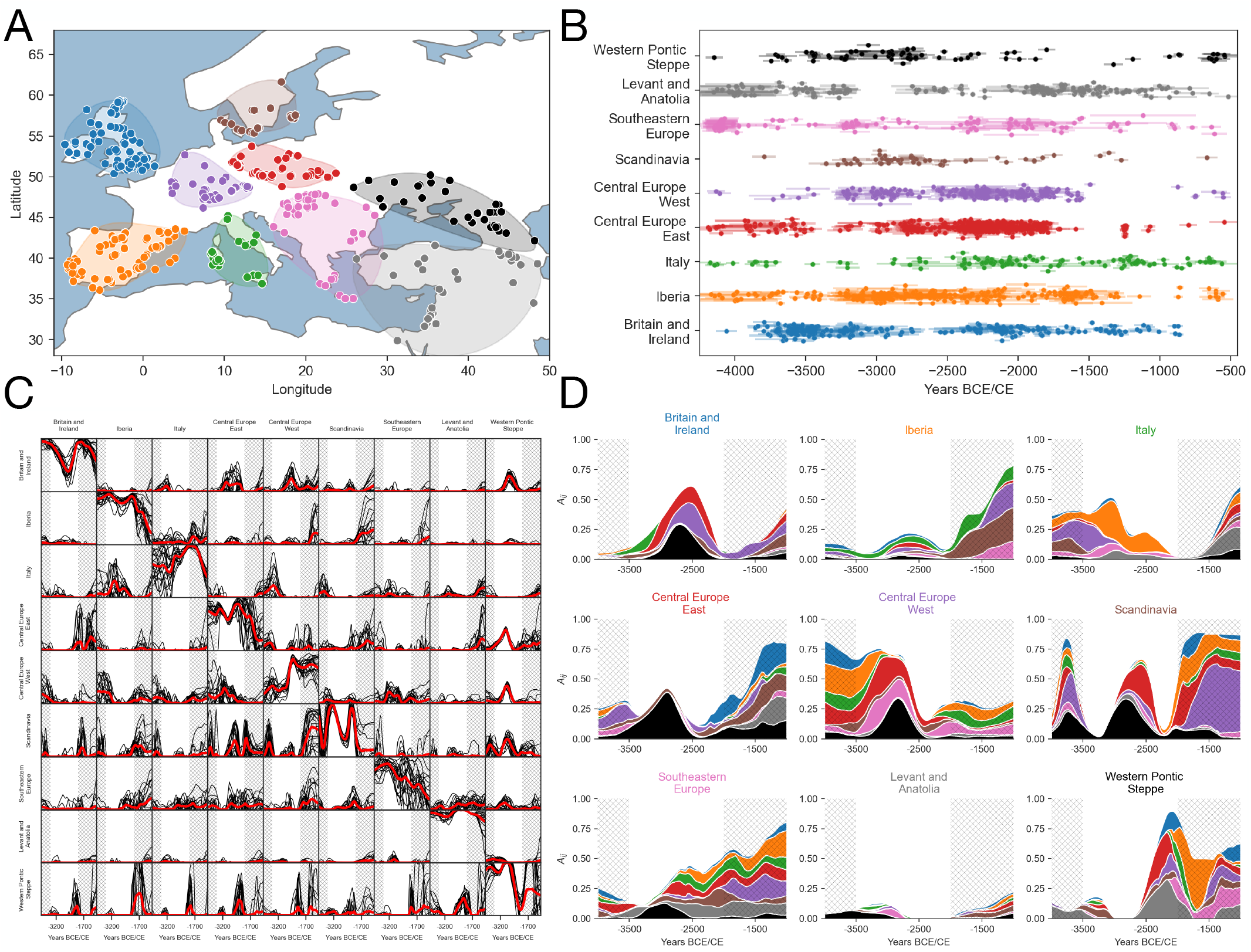
Time-resolved reconstruction of migration networks across nine regions. **(A–B)** Spatial and temporal distribution of the samples analyzed. **(C)** Time series for individual elements of the inferred migration matrix **A**, where **A**_*ij*_ denotes the fraction of genetic material that population *j* receives from population *i* within a 300-year interval. Black curves show interpolations from a single resampling iteration; the red curve represents the mean across iterations. Shaded areas indicate periods of lower confidence, based on a goodness-of-fit test (see main text and Supplementary Information, Fig. S7). **(D)** Each subplot shows the fraction of genetic material a focal region imports from all other regions over 300-year intervals. Titles indicate the focal region; colors indicate source regions. Values are means over resampling iterations (red line in subfigure C). The x-axis shows time.

To overcome the scarcity of samples in the time frame under examination, we implement an interpolation technique for both the *F*_2_ and 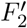 statistics, see the methods section for additional details and supplementary Fig. S6. We infer the migration matrix **A**(*t*) on the interpolated values throughout the entire time span. In Fig. 4C we summarise the results of our inference. We use a resampling approach to display the variability of the inferred matrix elements and use a goodness-of-fit test to characterize time periods where our method can predict the temporal dynamics of F-statistics at sufficient accuracy, see Methods and Supplementary Fig. S7. We observe that our method works well within the period of 3500 to 2000 BCE. Outside this period, the scarcity of samples, in particular for populations of the Western Pontic Steppe and Scandinavia, impedes an accurate interpolation. In Fig. 4D we visualize the average inferred matrix elements more intuitively. In each subplot, we visualize a single row *I* of **A**, which represents the proportion of migrations from all other populations *j* into population *i* through time.

In the time period of sufficient sample certainty, between 3500 and 2000 BCE, we find that Britain (blue) receives most of its genetic input from Central Europe (both East and West) and, by proxy, the Western Pontic Steppe. That is consistent with the well attested arrival of Steppe ancestry in Britain and Ireland in the third millennium BCE, together with the cultural markers of the Bell Beaker complex (10, 44, 45). Notably, the peak of Steppe-related inflow into Central Europe East occurs several hundred years earlier than the corresponding peak in Britain, consistent with a scenario in which migrating Steppe groups moved through Central Europe before reaching the North Atlantic coastline. By the time gene flow into Britain peaks, the inferred influx comes predominantly via Central Europe East and West, with only a modest direct contribution from the Western Pontic Steppe. This pattern suggests that the groups reaching Britain were often descendants of Steppe herders who had already acquired substantial ancestry from resident European farmer populations during their westward expansion.

Within our approach we can also integrate information across multiple time points to infer migratory events over extended periods. In Fig. S8-S9, we apply this coarse-graining method to the Steppe-related migration event and, once again, observe a clear east-to-west genetic flow; see also the SI text for additional information.

For Italy (green) we surprisingly observe relatively strong gene flow from Iberia around 3000 BCE, with smaller contributions from Southeastern Europe and Central Europe West. However, given few data points and our aggregation of regions with a relatively distinct genomic history (Sardinia, Sicily, Northern Italy), uncertainty is high in this case (see Fig. 4C). From published research we know that an influx of Steppe-related ancestry is only expected later in the second half of the third millennium (46), in Sicily then indeed possibly from Iberia (47). Steppe ancestry does not arrive in sizable proportions in Sardinia (48). Iberia (orange) itself receives little genetic input during this period, which is in line with previous research (49). Steppe-related ancestry appears there occasionally from 2500 BCE onward (45, 47) and becomes a stable component only by the end of the third millennium (50).

Central Europe East (red) shows strong genetic influx from the Western Pontic Steppe, consistent with the steppe migration event and the emergence of the Corded Ware cultural complex from 3000 BCE (51–53). Similarly and to no surprise, Central Europe West (violet) receives genetic input primarily from the Western Pontic Steppe, Central Europe East, and Southeastern Europe in the first half of the third millennium (8, 9, 54–56). The strong signal of ancestry flux from the West – from Britain, Italy and primarily Iberia – at around 3500 BCE is curious and potentially a methodological artifact. A similar signal has been observed already in (16), possibly explained by low levels of genomic differentiation among different Neolithic populations, creating a large geographic area of very similar genetic ancestry that is hard to resolve. Scandinavia (brown) shows contributions from both the Western Pontic Steppe and Central Europe East, most likely induced by the Steppe migration phenomenon before and after 2500 BCE (57, 58).

The result for Southeastern Europe (pink) is surprising in light of published research. The arrival of Steppe ancestry starts earlier here, and is a more gradual process compared to other parts of Europe (59–61). This may explain the weaker signal we observe from the East. The seemingly diverse influx from Northern, Western and Southern Europe throughout the third millennium, though, is hard to understand. Maybe this signal is driven by an elevated hunter-gatherer ancestry component with a potentially Central European source (62, 63). The Levant (grey) remains largely isolated in relation to the other regions considered here, with only a minor early contribution from the Western Pontic Steppe (61, 64– 66). Finally, the Western Pontic Steppe (black) itself appears to receive genetic input from all other populations toward the end of the study period, specifically from the South, so Anatolia, the Southern Caucasus or even the Levant. Due to sparse, spatially uneven sampling the error bars are large, indicating lower confidence in this new unexpected result (12, 61).

## Discussion

We presented a scalable and computationally efficient algorithm for inferring directional gene flow between populations. Its primary strength lies in its simplicity and minimal reliance on strong assumptions. Unlike methods that require the identification of ancestral populations or estimation of admixture proportions, our approach bypasses such prerequisites, making it broadly applicable and easy to implement. The algorithm’s scalability stems from its linear-time interpolation scheme and pairwise-statistic framework, which allow it to handle increasingly large ancient DNA datasets without prohibitive computational costs.

Simulation results demonstrated that the method reliably detects migration signals under realistic sampling constraints and data sparsity. We validated its effectiveness using two well-studied episodes of human history: the Early Neolithic expansion and the Western Steppe Migration. In the first case, we recovered asymmetrical gene flow from the Levant into Southeastern Europe after 7000 BCE, consistent with archaeological expectations. For the Western Steppe Migration, we demonstrated how combining information across multiple time points allows reconstruction of time-varying gene flow from the Pontic Steppe into Central and Western Europe during the third millennium BCE.

The second case also illustrates how secondary waves of migration can dominate the ancestry flux into a region: Britain appears to receive substantial Steppe-derived ancestry via Central Europe rather than directly from the Western Pontic Steppe. This is qualitatively in line with models of range expansions in which invading groups repeatedly pick up ancestry from resident populations along their route, so that later migrants are already strongly admixed by the time they reach the expansion front (67). Our framework makes such multistep, temporally structured ancestry flows directly visible in the time series of migration matrices.

Our current inference scheme relies on pairwise time bin comparisons, which may miss finer-grained temporal structure. We envision that extending the framework to a probabilistic time-series model — such as a Hidden Markov Model — could improve sensitivity and robustness by explicitly incorporating correlations across adjacent time points. This represents a promising avenue for future development.

A limitation of the current model is that it can only detect migration through its effect on reducing preexisting genetic divergence. As such, it cannot capture migration events that do not alter allele frequencies, such as exchanges between already well-mixed populations. Over longer time scales, and in contexts with high mutation rates, incorporating *de novo* mutations into the framework could expand its resolution and applicability. A more explicit incorporation of rare genetic variation may also provide an avenue for future improvement (68).

As genome sequencing efforts continue to improve spatial and temporal sampling density, we expect our method to become a valuable tool for reconstructing the dynamics of gene flow across broad regions and over extended historical periods.

Beyond estimating migration rates, our framework also provides a natural connection to classical models of admixture proportions. In the case where migration matrices **A**(*t*) vary over time, the cumulative effect of gene flow across multiple time steps is captured by the product **A**(*t*)**A**(*t* − 1)···**A**(1). Each row of this product matrix describes the expected proportion of ancestry in a given population at time *t* that originates from other populations at the initial time point. This product thus generalizes the notion of admixture proportions (8, 22, 69), allowing for temporally and spatially heterogeneous migration dynamics. In contrast to static models of ancestry composition, our rate-based formulation enables tracking the evolving structure and directionality of gene flow across both space and time.

## Methods

### Data preparation

For the empirical analysis we reused a dataset compiled for and described in (16), based on the Allen Ancient DNA Resource (AADR, V50.0, Dataverse V4.0, Oct 10 2021) (38, 39). This dataset includes samples from ancient human individuals genotyped at up to 1.23 million positions. Starting from the AADR we extracted samples with >25,000 recovered autosomal single-nucleotide polymorphism (SNPs), low contamination estimates, and favourable AADR quality assessment. Among closely related individuals or samples from the same individual we only kept the best sample. We removed SNPs in sections of the genome prone to linkage disequilibrium (70, 71) and below a minor allele frequency threshold of 5%.

For the spatial binning of samples into subpopulations, we defined spatial polygons based on Western-Eurasian geographical macro-regions. Their temporal binning was based on their archaeological dating information. Dating uncertainty was addressed with age resampling as in (16): If an ancient genomic sample came with one or multiple radiocarbon ages, then we drew age samples from the post-calibration distribution. If the dating information was limited to a range determined from contextual, archaeological information, then we drew samples from a uniform distribution within this range.

### The dynamics of neutral alleles in a metapopulation

Consider a population decomposed into *n* sub-populations, distinguished by location (different cities or districts) or any other feature, such as age or ethnicity. We assume to have available frequency time series of many unlinked alleles. Under neutrality, we can assume that the frequency *X*_*i*_(*t* + Δ*t*) of a particular neutral allele in population *i* at time *t* + 1 depends *linearly* on the allele frequencies {*X*_*j*_(*t*)}_*j*=1…*n*_ at some earlier time *t*,

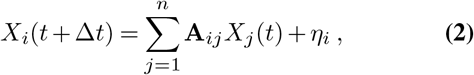

where **A** is the backward-migration matrix, i.e. **A**_*ij*_ is the proportion of individuals in population *i* at time *t* + 1 that originated from population *j* at time *t*. Thus, the elements are non-negative and, within each row, sum up to one, ∑_*j*_**A**_*ij*_ = *η*_*i*_ represents random genetic drift. For now, we do not need to know anything about the noise term, except that its expectation vanishes.

The linearity of Eq. 2 follows from the extensivity of the conditional expectation of the frequency of neutral alleles: Starting with half as many mutants at earlier time is expected to lead to half as many mutants at later time, i.e., 𝔼 [*X*_*i*_(*t* + Δ*t*)|*X*_*j*_(*t*) = *αx*_*j*_] = *α* 𝔼 [*X*_*i*_(*t* + Δ*t*)|*X*_*j*_(*t*) = *x*_*j*_] for any 0 *< α <* 1.

The rows of **A** have to sum up to 1 because neutral alleles are not expected to change if their frequency is the same in all populations, 𝔼 [*X*_*i*_(*t* +Δ*t*)|*X*_*j*_(*t*) = *x*] = *x* for any 0 *< x <* 1. Finally, negative matrix elements are excluded because they can generate negative expectations.

An alternative way of writing Eq. 2 is

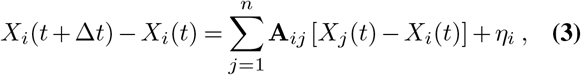

which explicitly shows that (i) the frequency in *j* only influences the frequency in *i* if *X*_*i*_ ≠ *X*_*j*_, and (ii) that a larger value of **A**_*ij*_ > 0 leads to a faster convergence of *X*_*i*_ to *X*_*j*_. The coefficient **A**_*ij*_ thus measures how population *j* influences population *i*.

### Expressing the dynamics in terms of *F*_2_ statistics

Our goal to infer **A**_*ij*_ can in principle be done directly using the dynamical equation for individual allele frequencies, Eq. 2, see also (18), via a state-space model approach that directly takes into account sampling noise, or via (total) least squares estimation by comparing predicted and observed lineage frequencies through time,

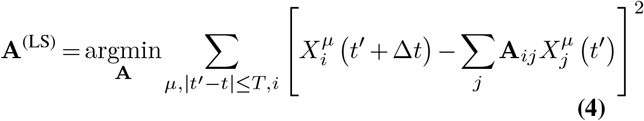

where the index *µ* labels the different genomic sites.

The downside of these direct approaches is that (I) it is very hard to reliably infer the frequency of individual alleles, (II) in the low sampling regime the noise is non-Gaussian and needs to be treated accordingly, and (III) one has to deal with complicated biases introduced by which set of genomes actually have a given allele. While the last point can perhaps be avoided by focusing on the set of overlapping alleles present in all genomes, this would reduce the amount of data that can be used, especially in ancient DNA.

Here, we take a simpler approach and derive an equation that constrains the mean allele frequency differences between different populations at different times. This leads to *F*_2_ statistics and a dynamical extension thereof, which can be robustly inferred from the data.

For notational simplicity, we introduce the following abbreviations: 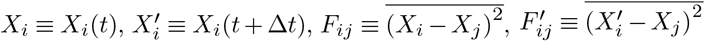, where 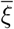 refers to the combination of taking the expectation value of *ξ* and averaging over all biallelic sites in a genome. In practice, if we have many sites, averaging over all alleles should generate a narrow Gaussian around the expectation value. Note that *F*_*ij*_ = *F*_2_(*X*_*i*_, *X*_*j*_) is the standard *F*_2_ statistics (22), which corresponds to the Euclidean mean squared length of the allele frequency vectors of populations *i* and *j* normalized by the number of alleles. The primed quantity *F* ^*′*^ corresponds to the time-dependent version of this statistics, i.e., it is the normalized Euclidean mean squared length of the allele frequency vectors of population *i* at time *τ* and population *j* at time *t*.

### Derivation of the equations of motion in terms of F_2_ statistics

We derive 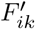 for two different populations, which will give us a simple linear equation that is well suited for the inference of **A**. We begin with the simple identity

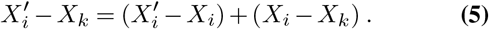

Taking the square of both sides and averaging over all alleles, we obtain

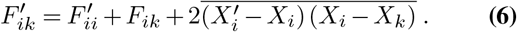

The average on the right hand side can be rewritten using our basic equation of motion Eq. 3 as

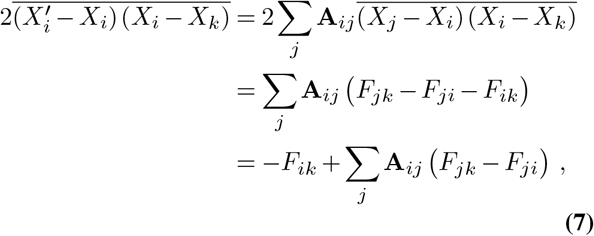

where in the final step we used the row normalization Σ_*j*_ **A**_*ij*_ = 1 and in the second-to-last step we used the identity:

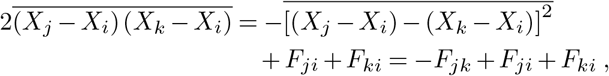

which corresponds to the statistics commonly known as *F*_3_(*X*_*i*_; *X*_*j*_, *X*_*k*_). Inserting Eq. 7 into Eq. 6, we obtain the useful linear equation

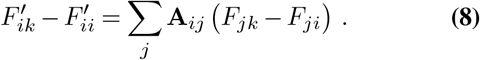

This is a set of *n* × (*n* − 1) equations to fix the *n* × (*n* − 1) components of the **A** matrix, given we can observe all *F* ^*′*^ and all *F* . Remarkably, these equations do not contain a term from genetic drift, which cancels on the left hand side in the difference of two *F* ^*′*^ statistics. Note that, while *F*_*ij*_ is symmetric in its indices, 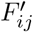 is not because of the time delay.

### Estimation of F2 statistics from data

Following (22) we compute an unbiased estimator for the *F*_2_ statistics under the assumption of binomial noise at a single site between populations A and B as:

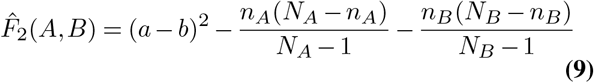

where *n*_*K*_ are the allele counts in the population *K, N*_*K*_ is the total counts of both alleles, *k* = *n*_*K*_*/N*_*K*_ is their frequency. In order to produce confidence intervals in the inference of **A**_*ij*_ we compute the average of 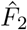 (*A, B*) in blocks of nearby SNPs (typically 5000) and we bootstrap blocks.

### Inference of the migration matrix A via convex optimization

The linear equation Eq. 8 suggests a straight forward approach to infer **A**, namely via linear regression under the constraints **A**_*ij*_ > 0, Σ_*j*_ **A**_*ij*_ = 1. This will give us an MLE estimate of **A** since all *F*’s should be approximately Gaussian.

Minimizing the square difference of left and right hand sides of Eq. 8 for all *i* and *k* reduces to the following optimization problem

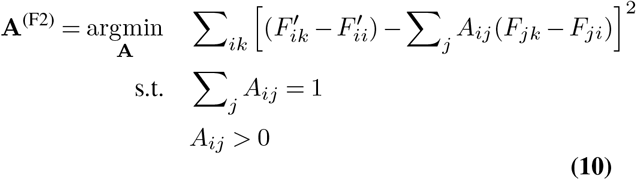

We rewrite this constrained least-squares optimization problem to emphasize that each row of the matrix **A** ∈ ℝ*n*×*n* can be optimized independently. Define the row vector 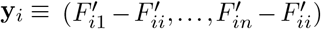 and the matrix **X**(*i*) with elements 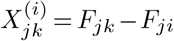. Then, each row vector **a**_*i*_ ≡ (*A*_*i*1_, …, *A*_*in*_) of the matrix **A** can be estimated independently via constrained linear regression,

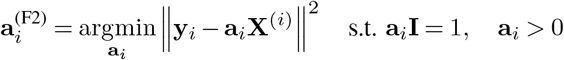

This is a standard non-negative least-squares problem with a simplex constraint, and can be solved efficiently and independently for each row (**a**_*i*_)_*i*=1,…,*n*_.

### Estimation of Confidence Intervals

We estimate confidence intervals via multiple iterations of resampling of our data with respect to different sources of potential uncertainty: (I)individual variability, (II) dating uncertainty, (III) genetic uncertainty. For each sample, we have a metapopulation assignment and its date. We first define a binning Δ*t* and a period of interest **p** = (*T*_0_, *T*_0_ + Δ*t, T*_0_ + 2Δ*t*,… ) for the analysis. As there is an underlying uncertainty in dating, we seek to take it into account via temporal resampling following the same approach of (16). The main inference algorithm is subdivided into the following steps:

- Time resampling of dates.
- Subsampling of *N* − 1 individuals from the pool of *N* available samples in each time bin and metapopulation. If only 2 individuals are available, keep both.
- Assignment of samples to bins in **p** and compute count vector **n**_*t*_ and total count vector **N**_*t*_ for each time bin.
- Computation of mean *F* 2 and *F* 2*′* statistics via block-wise resampling.
- Infer migration matrix **A** via the constrained convex optimization problem as described in eq. 9.

By iteratively following the described process several times, we can compute confidence intervals for the migration matrix elements **A**_*ij*_ that take into account the sampling of individuals, time uncertainty in the samples dating, and uncertainty in the computation of F2 summary statistics.

### Interpolation Analysis

The interpolation analysis algorithm comprises several steps, which are delineated as follows: (I) Sampling multiple *F* and *F* ^*′*^ values across the entire period via temporal resampling. (II) Interpolating using a convolution filter. For the initial sampling step (I), we do the following:

- Randomly sample the start of the first time bin from the uniform distribution 𝒰 (−4150, −3850).
- Sample dates for all individuals and allocate them into corresponding bins.
- Calculate *F* and *F* ^*′*^ for each pair of populations across all time bins.

By iteratively conducting the aforementioned process, two sets of tuples, denoted as *D*_*F*_ = (*t, F* ) and *D*_*F*_ *′* = (*t, F* ^*′*^), are constructed, where *t* represents the start time of the time bin. In the subsequent step, resampling and interpolation of the points from *D*_*F*_ and *D*_*F*_ *′* are performed to assess the robustness of the interpolation line. The interpolation algorithm (II) entails the following steps:

- Define dense binning of size 30*y* across the period.
- Sample *N* = 50 tuples from *D*_*F*_ and *D*_*F*_ *′* and allocate the values to the bins.
- Compute the average within each bin and infer the value from neighboring bins if no value is present within the bin.
- Perform multiple (N=5) convolutions with a uniform filter of size of 10 bins to perform smoothing.

Although using convolutions for smoothing can be susceptible to edge issues, we can mitigate this problem by extending the time period under consideration. While more advanced interpolation techniques could be applied, such as Gaussian processes or Kalman smoothing, we leave this for future research and development.

### Goodness-of-fit test

The inference of migration matrix elements is carried out one row at a time. For each row, the model’s performance can be assessed by calculating the Pearson correlation between the left and right-hand sides of Eq. 2 using the inferred matrix. In Fig. S7A, we conduct this goodness-of-fit test across the entire time span for each row of **A**. We observe that, especially for the Western Pontic Steppe and Scandinavia metapopulations, there is a period where the method shows reduced accuracy. In Figure S7B, we plot the left and right-hand sides of Eq. 2 during the period of high accuracy (3500 - 2000 BCE) to provide a clearer visualization of the model’s predictions.

### Lower Dimensional Representation

The representation of genetic samples into lower dimensions to better visualize genetic relationships is typically performed via PCA projection. This procedure is equivalent to Multidimensional Scaling on the matrix of all pairwise F-statistics (25). While this procedure is typically performed independently of temporal sampling information, it is possible to extend this idea to include time. At each time point *t*, we perform centering of the matrix **M**(*t*) = −0.5**CF**(*t*)**C** with *C* = **I** − (1*/n*)**1** is a centering matrix and **I, 1** the identity matrix and a matrix of ones. We calculate the average of the first two eigenvectors of **M**(*t*) over time and then project all **M**(*t*) into a two-dimensional space. In Fig. S7C-D, we present the resulting lower-dimensional representation of our populations. As the eigenvectors and eigenvalues of **M**(*t*) evolve over time, as shown in Supplementary Fig. S7E-F-G, we opted to compute the average of the first two eigenvectors across time to determine a consistent projection for all **M**(*t*). This method yields a lower-dimensional representation of the populations’ relative genetic trajectories. Populations with a higher genetic exchange between them are brought closer together, while those that remain genetically isolated move further apart.

## Code and Data Availability

All code and data to reproduce the results of this paper can be found at 10.5281/zenodo.18740249 and https://github.com/Hallatscheklab/interactions-from-fluctuations/

## Acknowledgements

We thank all members of the Hallatschek lab (past and present) for helpful discussions and advice on the project. This work was supported by the National Institute of General Medical Sciences of the NIH under award R01GM149827 and by a Humboldt Professorship of the Alexander von Humboldt Foundation. TO acknowledges support from JSPS KAKENHI (Grant Numbers JP22K03453, JP22K06347, 25K09723, and 25K07168) and the RIKEN iTHEMS Program. GI acknowledges support from Humboldt Research Fellowship.

## Supplementary Information

### Wright-Fisher Simulations

Our Wright-Fisher simulations between *N*_*D*_ metapopulations are defined by the following steps:

- **Initialization:** once at beginning. Initially, allele frequency distributions are drawn via

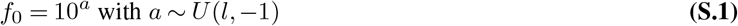

where *U* (*l*, −1) represents a uniform distribution in the interval [*l*, −1] with *l* a parameter. We generate *N*_traj_ distinct lineages for each population *i* we sample

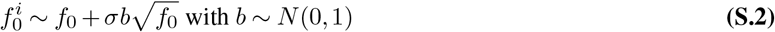

where *σ* is another parameter that modulates the initial genetic distance between the metapopulations.
- **Generation:** for T times. At each new generation *t*, we first determine the updated allele frequencies by calculating

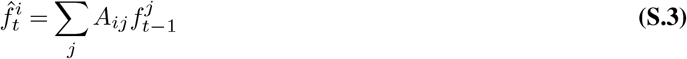

where the matrix **A**_*ij*_ dictates how individuals are redistributed among metapopulations and *f*_*t*_ values are constrained in the [0, 1] interval. We then draw *N*_pop_ samples via a binomial distribution

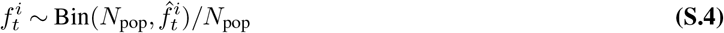

for each metapopulations and lineage. Observations are performed via an additional binomial sampling 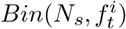 at each generation, which depends on the parameter *N*_*s*_.

### Benchmark of inference method on simulated data

This section aims to evaluate and compare the inference accuracy of our F2-based estimator (Eq.8 in the main text) with that of a straightforward estimator derived from least squares regression (Eq.4 in the main text).

We evaluate the effectiveness of our methods using Wright-Fisher simulations across several metapopulations. Our objective is to investigate how various simulation parameters affect the performance of the methods.

#### Key Parameters for Comparative Analysis

To evaluate the complexity of the task, the following parameters are varied in the benchmark:

- *l* defines the range of initial frequency magnitudes; at decreasing frequencies, the sampling noise increases and the complexity of the inference as well.
- *σ* influences the initial genetic distance among metapopulations; larger genetic distances lead to easier inference of **A**_*ij*_.
- *N*_*s*_ determines the extent of sampling from the metapopulation, which is related to the level of observational noise; the higher the observational noise, the harder the inference.
- *T* represents the entire duration of the mixing process; a longer observation period provides more extractable information.
- *N*_*D*_ refers to the number of metapopulations; Since a squared matrix is being inferred, the number of parameters increases quadratically with *N*_*D*_.

#### Evaluation

The default parameters of the simulation are: *σ* = 0.1, *T* = 5, *l* = −1, *N*_*D*_ = 5, *N*_traj_ = 5000, *N*_pop_ = 1000 and *N*_*S*_ = 100. We compute the mean squared error between the ground truth values **A**_*T*_ and the inferred values from the least square method **A**_*LS*_ and the F2-based method **A**_*F* 2_ on multiple independent simulations at varying parameters. The results of the analysis are depicted in Fig. S3 . The *F* 2 method consistently outperforms the naive *LS* method across all parameter regimes.

### Zooming into the third millennium BC

In this section, we demonstrate the ability of our method to integrate information from more time points to infer migratory events across extended periods of time. We focus on the third millennium BCE, a period where many studies have uncovered substantial migration of populations from the steppe regions north and east of the Black sea into both Western and Eastern Europe (8, 9), as also reflected in our extensive analysis of Fig. 4.

We conducted an analysis of samples from predefined geographic locations dating back to -3300 and -1800 BCE, employing time intervals of 300-year bins. These samples consist of a subset of the samples analyzed for Fig. 4, where we exclude Iberia, Italy, and Scandinavia, as their total average input genetic flux within that time period is only ∼ 7% for those populations. We initially quantify gene flow between each pair of consecutive time bins (Fig. S8A). Similarly to the analysis in the previous section, we observe here that the two Central European metapopulations experience significant gene flow from the Western Pontic Steppe in the first half of the observed period. The Britain and Ireland metapopulation experiences substantial gene flow from the two Central European populations in the second half of the observed period. The Western Pontic Steppe primarily acts as a source of gene flow.

We seek then to integrate information across all time bins and infer a coarse-grained migration matrix that summarizes the average gene flow over the period of interest. This can be achieved by adjusting the minimization equation in 10, averaging over all time points, and inferring a single migration matrix for all times. We additionally implement a simple constraint on neighbouring edges during inference in order to avoid serial passaging (A->B->C) mimicking long-range direct connections (A->C), see Fig. S8B.

In Fig. S8C, we present the inferred migration matrix **A**_*ij*_ resulting from this procedure, with populations ordered from West to East. We observe that the values in the upper triangular part are significantly higher than those in the lower triangular part, emphasizing a prevalent east-to-west migration trend. Moreover, the strongest elements are typically on the diagonal, consistent with the idea that the main source of genetic composition for a population in the next time step is the population itself at the previous time. The exceptions are Central Europe West and the British Isles, both of which have the highest migration coefficients from Eastern Central Europe and Western Central Europe respectively.

As a robustness test, if we further merge the two Central European populations into a single group and repeat the analysis, we still observe a pronounced east-to-west genetic flux, see Fig. S9.

In Fig. S8D we depict the strongest off-diagonal elements of the matrix (thresholded at 0.1 of the average value) to offer a clearer visual representation of these population interactions on the map. This representation provides enhanced clarity regarding the aforementioned points. In Fig. S8E-F we present a visualization of the matrix’s spectral decomposition.

**Fig. S1.**
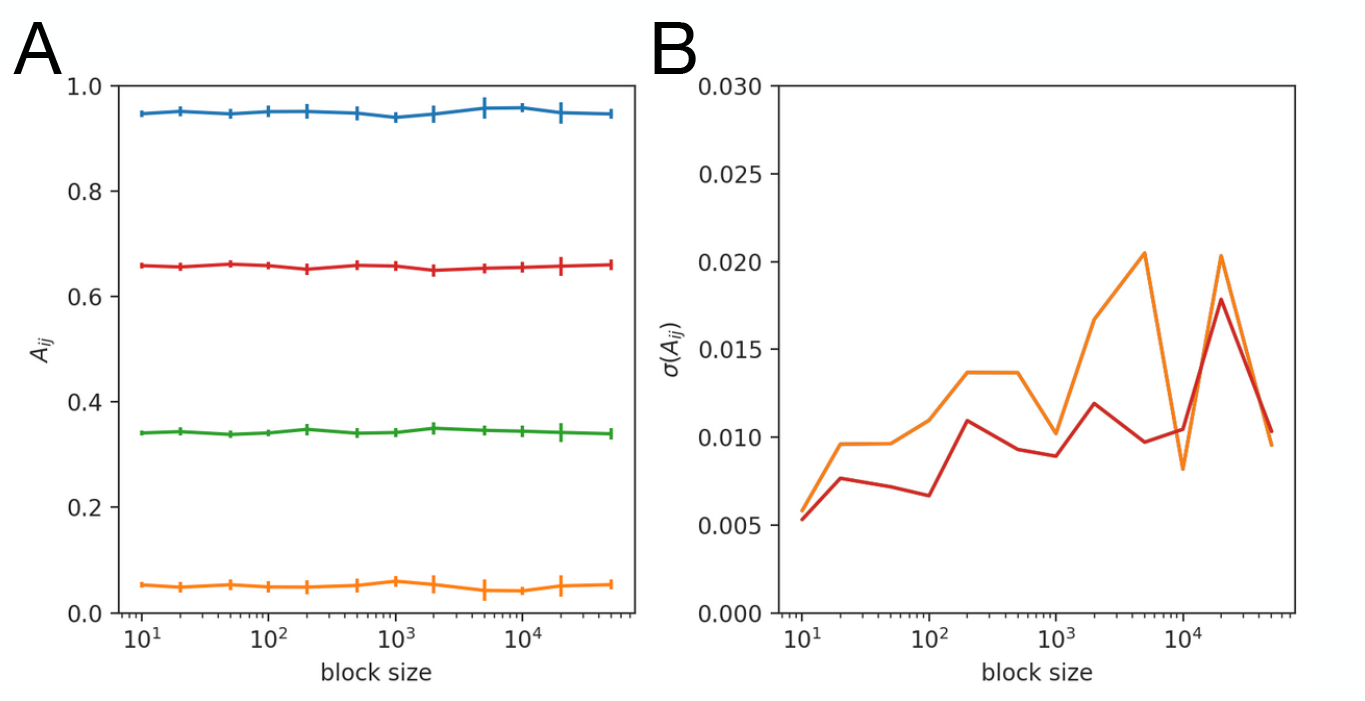
**(A)** Mean and standard deviation of migration matrix elements computed via resampling of genetic blocks of increasing size, each color identifies a different element of **A**. The evaluation is done on the same samples of Fig. 3 at the 4th time bin. **(B)** Standard deviation in the estimation of **A** as a function of block size. We find a limited dependency on block size.

**Fig. S2.**
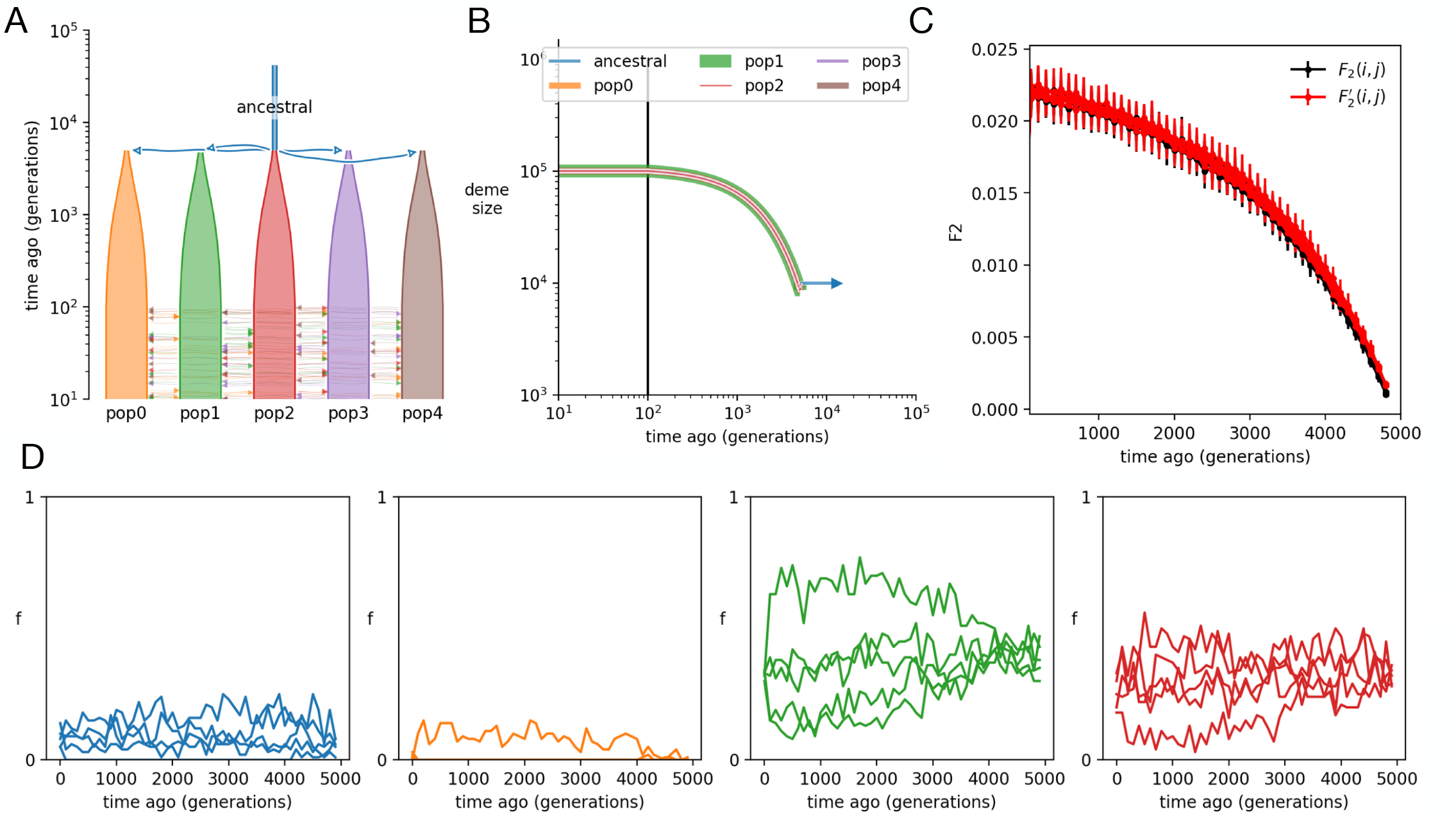
Establishment of genetic diversity across populations in the simulations. **(A-B)** We simulate a similar scenario to that of Fig. 2 but sample across the whole period of isolation and expansion of the populations. **(C)** During this period the genetic divergence between populations monotonically increases. **(D)** Examples of lineage trajectories over the sampled period.

**Fig. S3.**
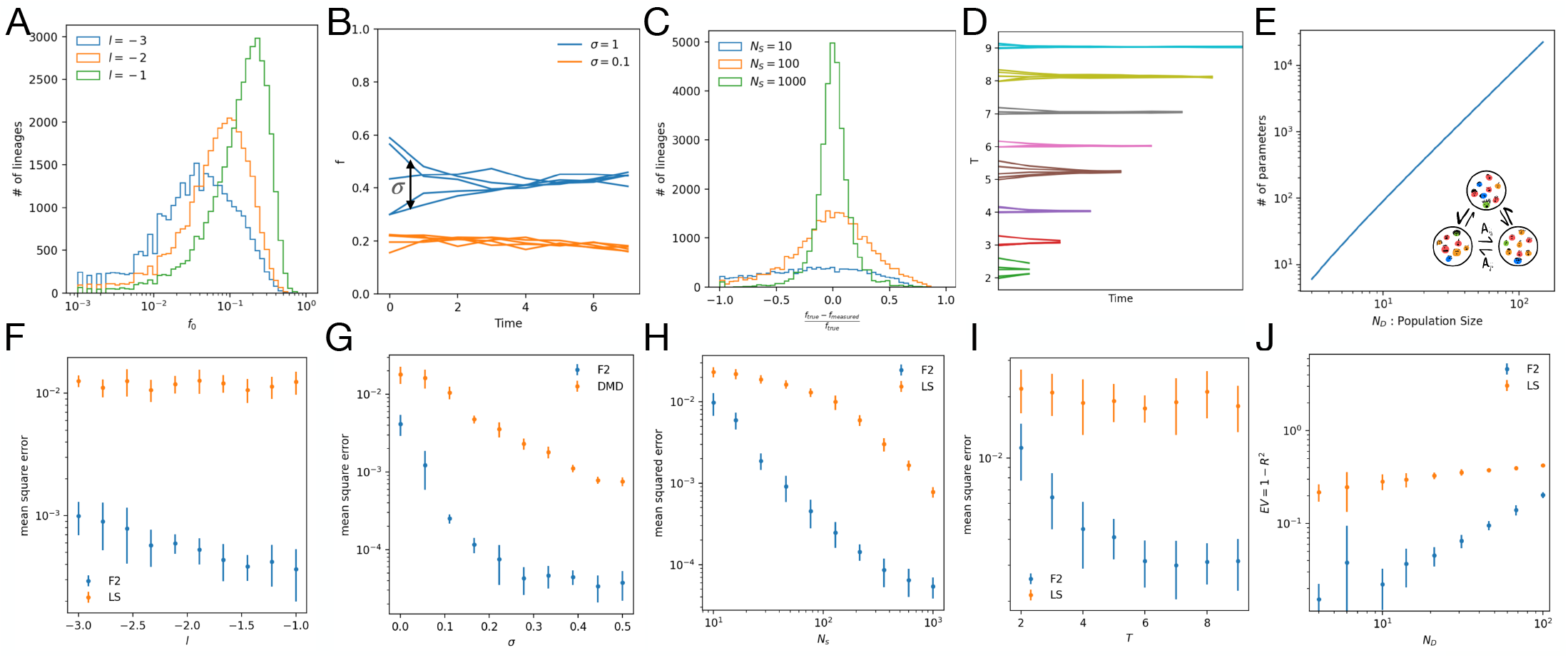
Benchmark on simulated data of our F2-based method and least squares estimation (LS) We perform Wright Fisher simulations of multiple metapopulations where at each generation **A**_*ij*_ is used to reassign individuals between populations, see SI text for more details. **(A)** Distribution of initial allele frequencies 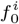 for different values of *l* and **(B)** depiction of lineage frequency trajectories as a function of the variance in the initial allele frequencies across demes *σ*. **(C)** Distribution of error induced by sampling noise as a function of *N*_*S*_ and **(D)** examples of different lineage trajectories for increasing *T* . **(E)** Given *N*_*D*_ populations, (*N*_*D*_ − 1)^2^ parameters need to be inferred. **(F)** As expected, for bigger values of l, which induce a larger fraction of more common alleles, the inference is more accurate. **(G)** Our inference relies on detecting changes in allele frequencies through time. Thus, if *σ* is bigger, these differences are more significant and the inference is easier. **(H)** The more individuals are sampled at each generation, i.e. lower observational noise, the better the accuracy of the inference. **(I)**The method can integrate information across multiple time steps. We find an increase in accuracy as a function of the number of time steps until saturation. **(J)** As the number of parameters to be inferred increases, the accuracy is degraded. In this last benchmark we introduce increasingly high sparsity in the matrix when increasing the number of metapopulations to have a reasonable comparison across different parameter regimes. We estimate the error via the fraction of unexplained variance of non-zero matrix elements *EV* = 1 − *R*^2^.

**Fig. S4.**
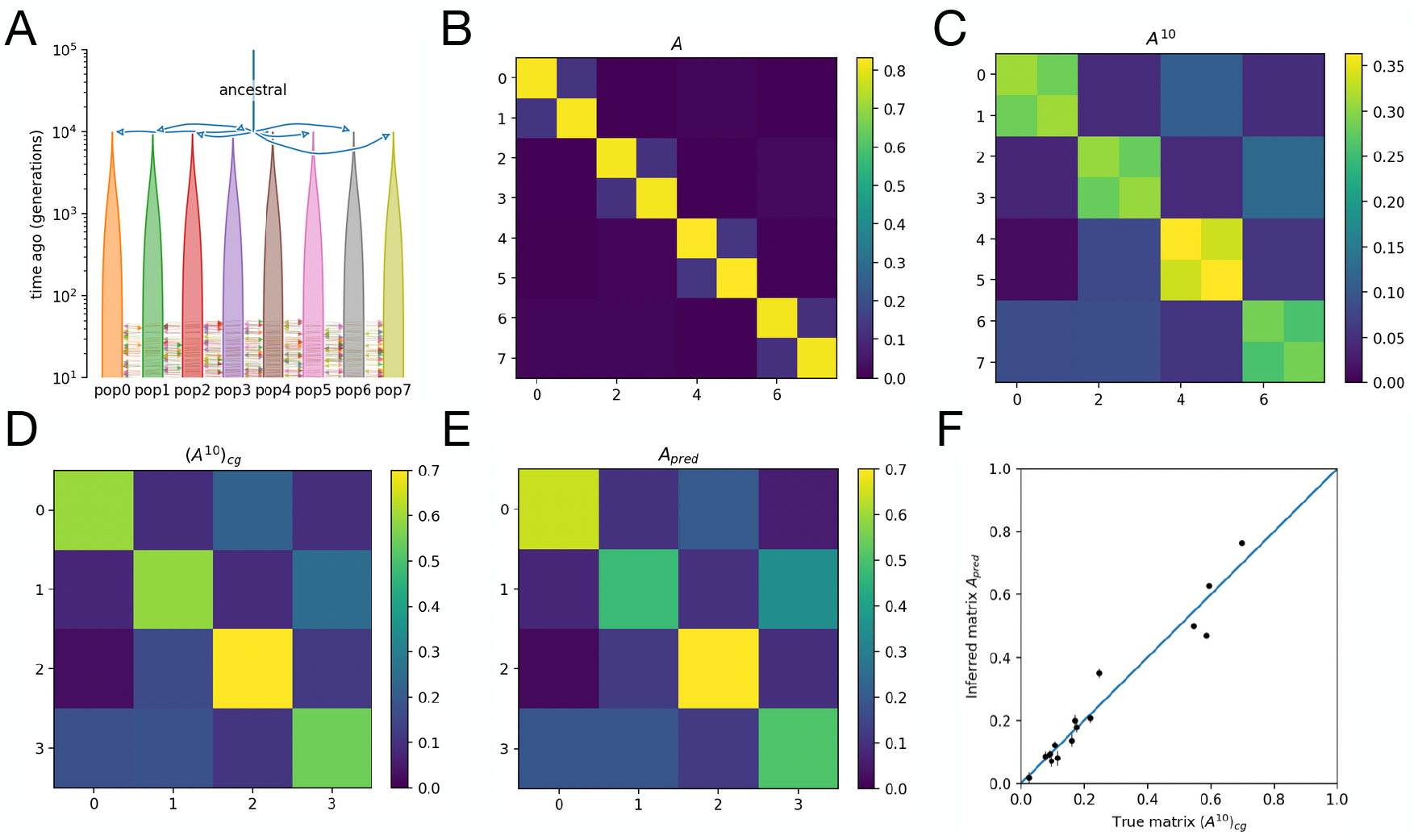
Stability of inference when the meta-population has a substructure. **(A)** We repeat the simulation following the same parameter of Fig. 2 with 8 populations that are organized in pairs of more strongly connected demes. **(B)** Migration matrix used for the simulation **(C)** Since we resolve the dynamics every 10 generations, the time traces follow **A**^10^ . **(D)** Coarse graining of the matrix is performed by: (I) a weighted average over the rows where the weights are proportional to the lineage steady-state distribution; (II) the sum of the elements of the resulting vector. **(E)** Inferred coarse grained matrix **(F)** Scatter plot of matrix elements of the true and inferred coarse grained matrices.

**Fig. S5.**
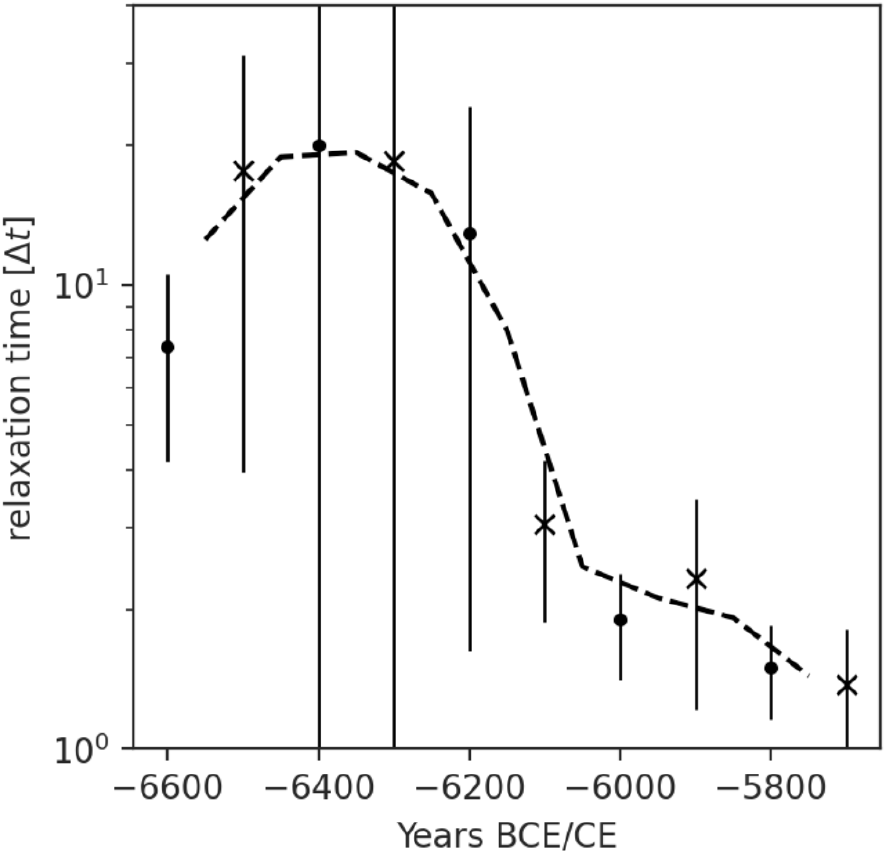
Spectral decomposition and relaxation times. Estimation of relaxation time *t* = (1 − *λ*_2_ )^−1^ by eigenvalue decomposition *λ*_2_ of the inferred migration matrices of Fig. 3. For each time slice, eigenvalues *λ*_*i*_ were computed via eigendecomposition of the migration matrix *A*. The dominant eigenvalue *λ*_1_ = 1 reflects conservation of probability mass, while the subdominant eigenvalue *λ*_2_ determines the asymptotic rate of convergence toward the stationary distribution. Error bars represent variability across bootstrap realizations of the inferred migration matrices.

**Fig. S6.**
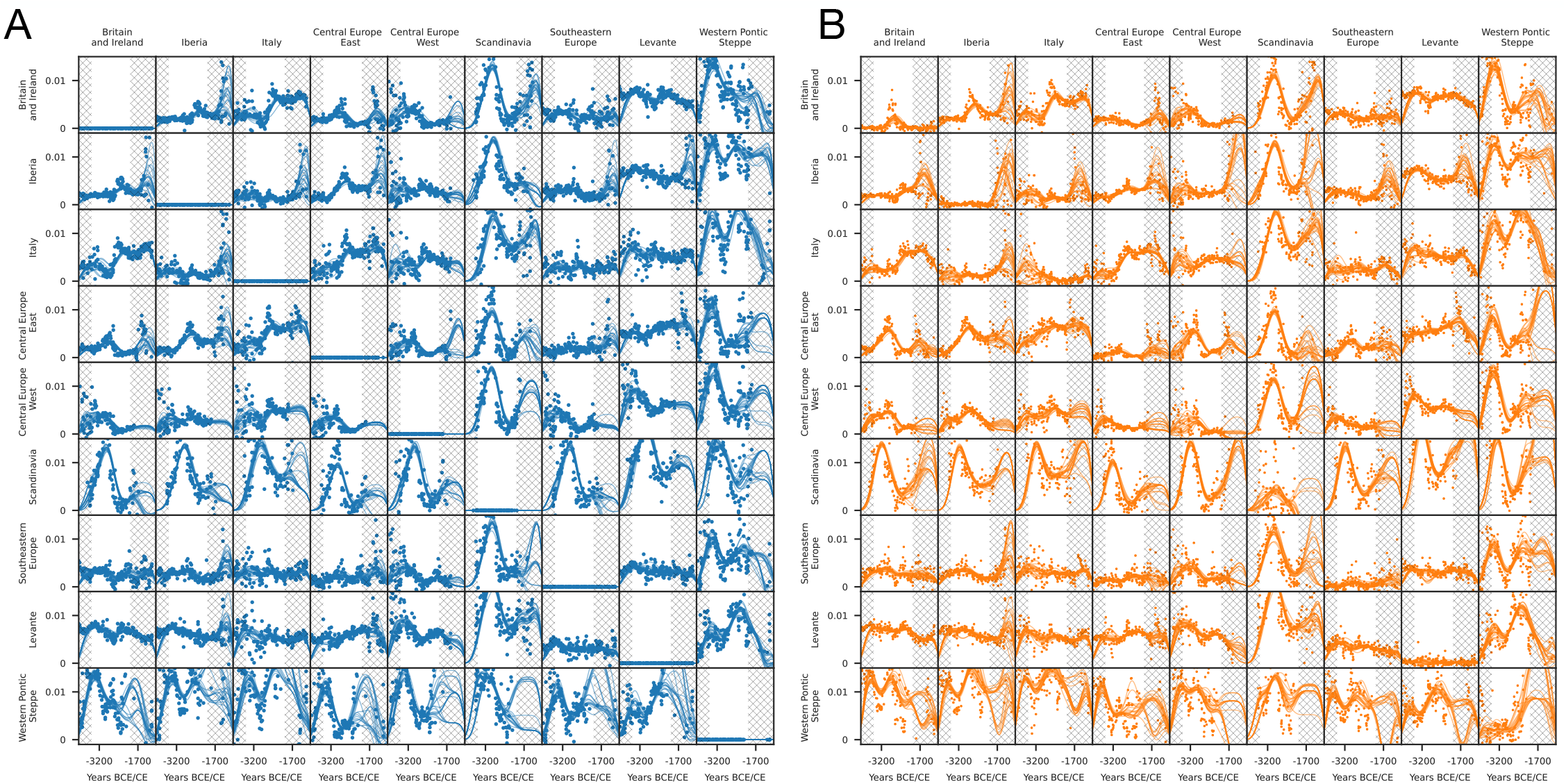
Large Scale Analysis with Interpolation. **(A)** Matrix of all pairwise comparisons of *F* statistics across time with interpolation lines. **(B)** Matrix of all pairwise comparisons of time-lagged *F* ^*′*^ statistics across time with interpolation lines. Crossed area refers to a region with lower confidence in our model. See Fig. 4 and Methods for more details.

**Fig. S7.**
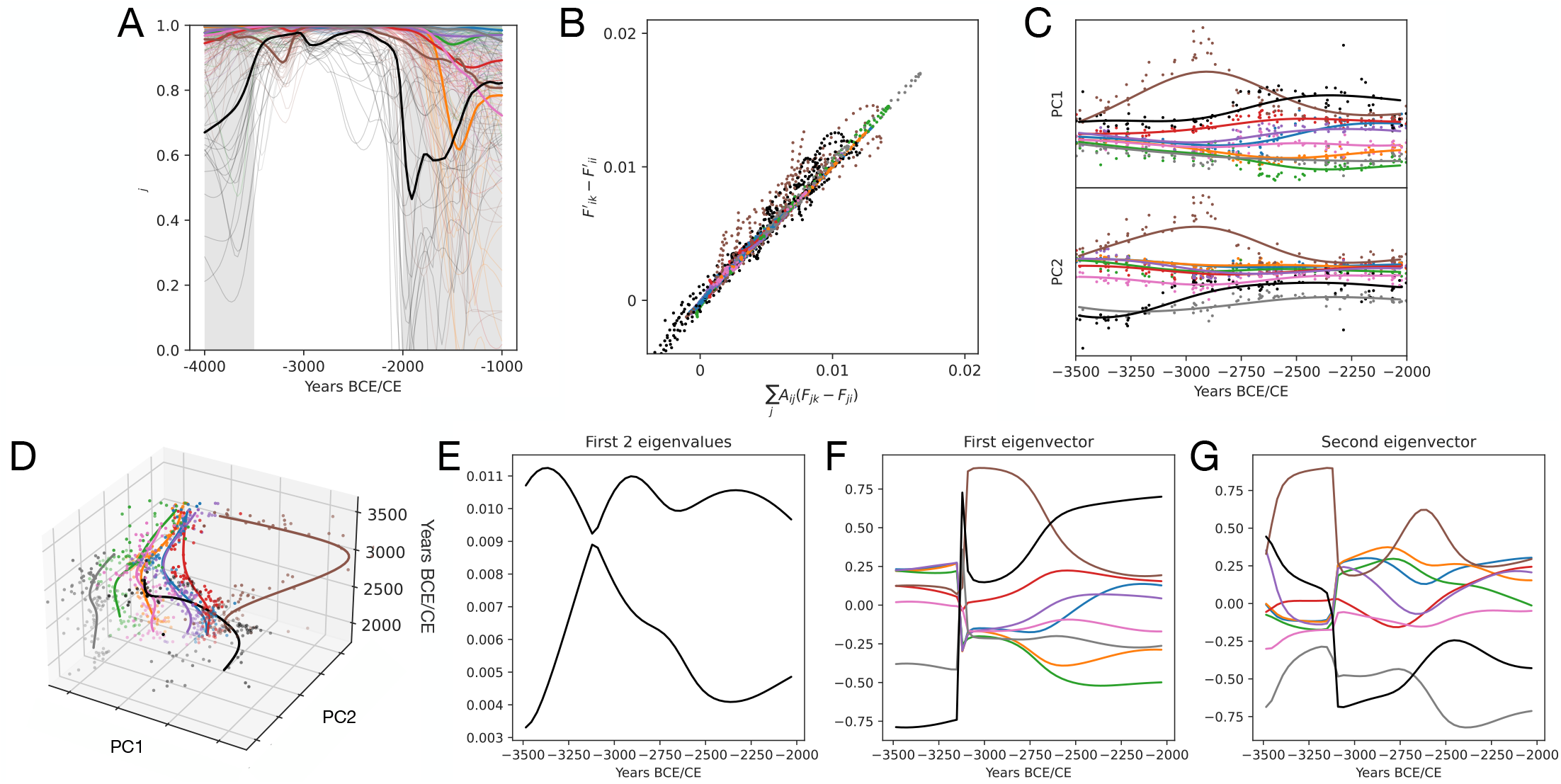
Goodness-of-fit and Lower Dimensional Representation. **(A)** Pearson correlation of left and right-hand side of Eq.2 for each row of **A** to check the consistency of the model. **(B)** Plot of left and right-hand side of Eq.2 using the inferred migration matrix **A** within the period of high confidence (3500 - 2000 BCE). Western Pontic Steppe (black points) and Scandinavia (brown points) are the populations with lower confidence and higher errors. **(C-D)** Lower dimensional mapping of the F and F’ matrices across time with average interpolation lines. The mapping is defined by the average first 2 eigenvectors of the centered F matrix across time *M* = −0.5*CF*_2_ *C*, see Methods for additional details. **(E)** First 2 eigenvalues of *M* across time. **(F)** Elements of the first eigenvector of *M* across time. **(G)** Elements of the second eigenvector of M across time.

**Fig. S8.**
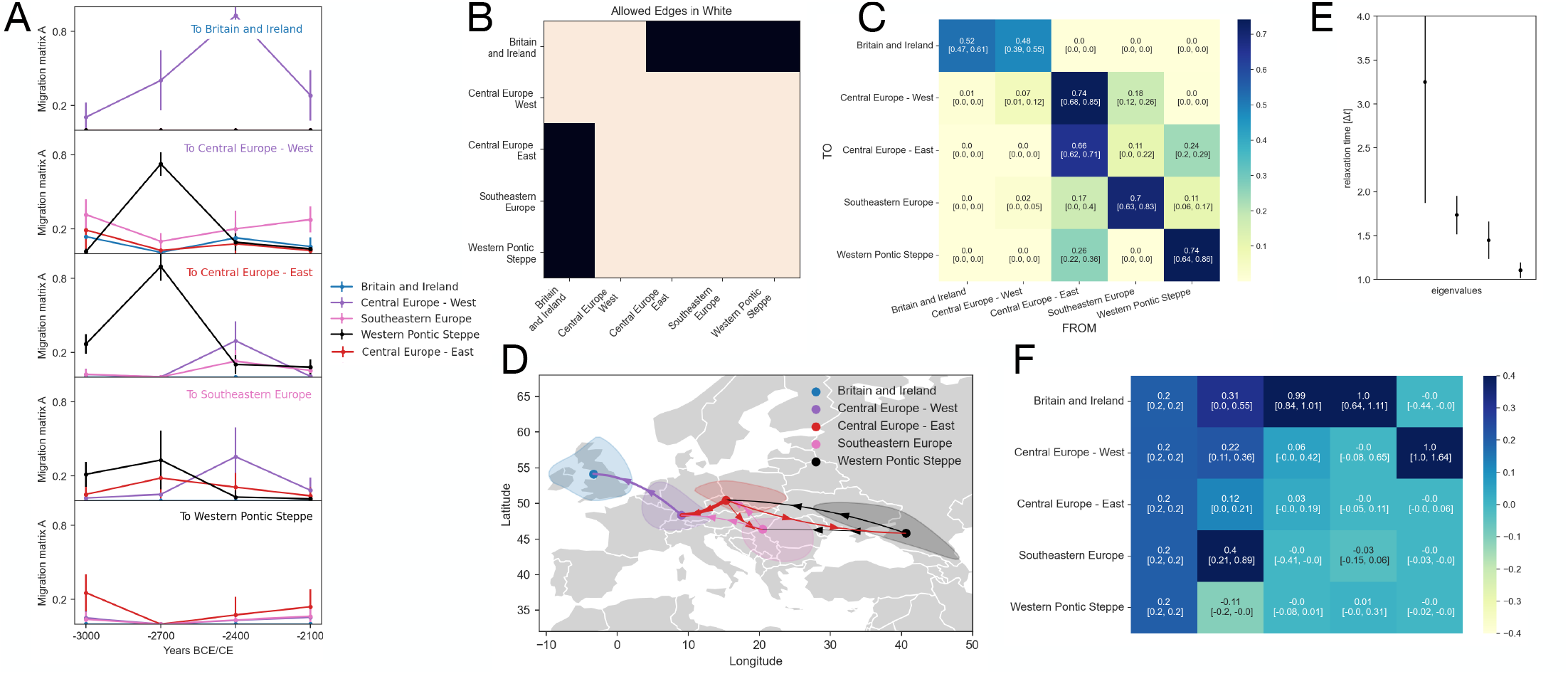
Onset and dynamics of the steppe migrations. We seek to analyze the migration of Yamnaya herders from the Western Pontic Steppe into Europe, beginning around 3000 BCE. **(A)** Inferred migration over the entire time period, with each plot showing one row of the migration matrix **A**(*t*) at each time point. **(B)** Allowed edges during migration inference at the coarse-grained level. **(C)** Inferred migration after coarse-graining across time windows. For each matrix element, the mean value is shown along with the lower and upper quartiles. The pronounced upper-diagonal structure reflects westward movement, as populations are ordered from east to west in the matrix rows and columns. Confidence intervals correspond to the quartiles. **(D)** Visualization of the strongest migration links (values > 0.1) as directed arrows.**(E)** Relaxation times derived from the eigenvalues of the inferred matrix **(F)** Right eigenvectors from spectral decomposition of the inferred matrix.The first eigenvector identifies the steady state probability of the system and has the corresponding eigenvalue *λ* = 1.

**Fig. S9.**
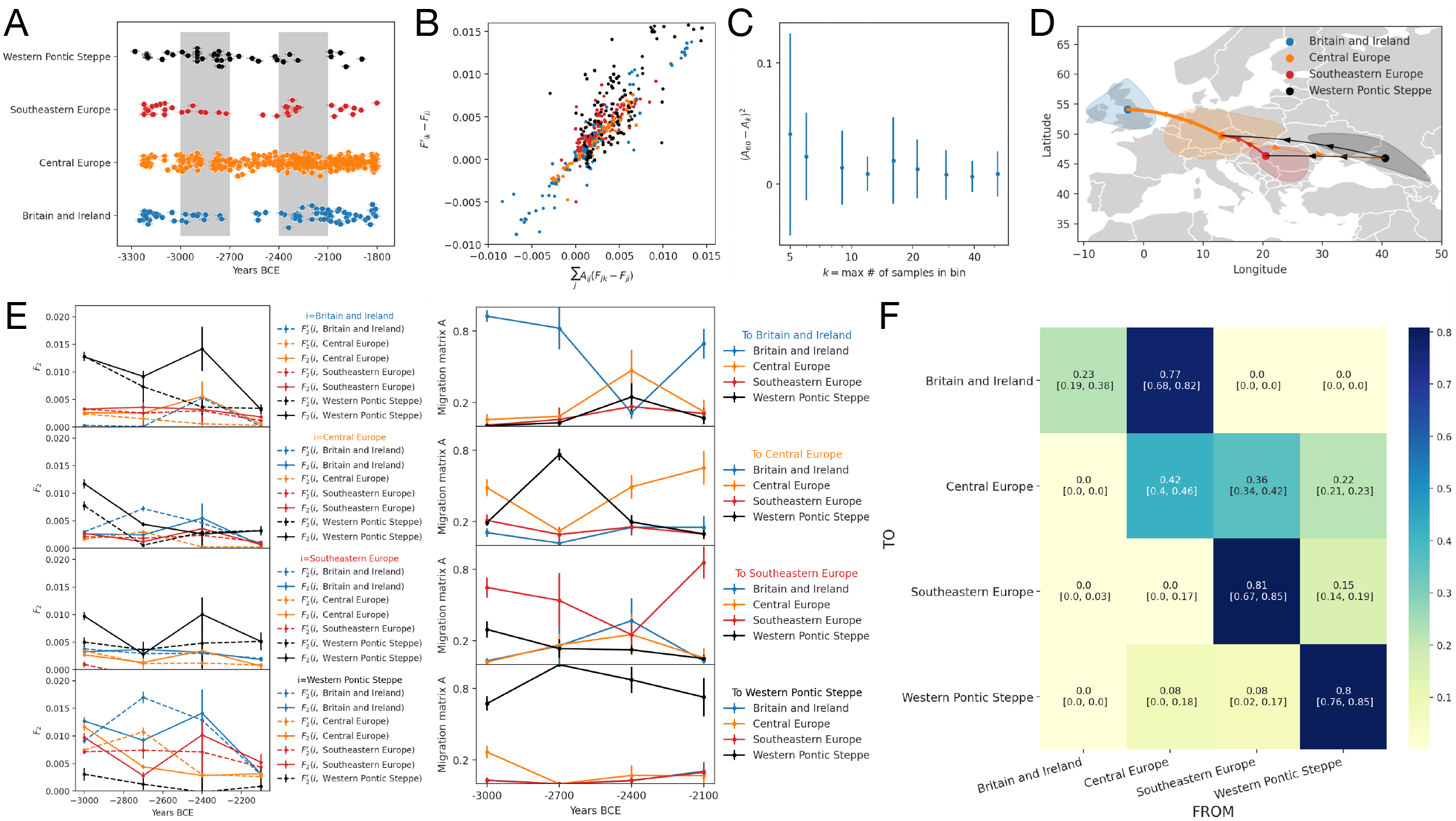
Coarse graining of the two Central European clusters. Same analysis as in Fig.S8 but with “Central Europe - West” and “Central Europe - East” combined into one cluster. **(A)** Temporal organization of the samples analyzed. **(B)** Comparison of model prediction ( the left side of Eq. 1) and observed time-lagged F2 statistics (the right side of Eq. 1) using the inferred matrix. **(C)** Stability of matrix inference with respect to subsampling of individuals in the Central Europe population within each bin. **(D)** Visualization of the strongest matrix elements define by the average matrix value bigger than 0.1. **(E)** Temporal variation of *F*_2_ statistics (left) and inferred matrix elements (right) across the analyzed time period. **(F)** Inferred migration matrix obtained by integrating information across the whole time period.

